# Identifying Potential Causal Risk Factors for Self-Harm: A Polygenic Risk Scoring and Mendelian Randomisation Approach

**DOI:** 10.1101/673053

**Authors:** Kai Xiang Lim, Frühling Rijsdijk, Saskia P. Hagenaars, Adam Socrates, Shing Wan Choi, Jonathan R.I. Coleman, Kylie P. Glanville, Cathryn M. Lewis, Jean-Baptiste Pingault

## Abstract

**Background:** Multiple individual vulnerabilities and traits are phenotypically associated with suicidal and non-suicidal self-harm. However, associations between these risk factors and self-harm are subject to confounding. We implemented genetically informed methods to better identify individual risk factors for self-harm.

**Methods:** Using genotype data and online Mental Health Questionnaire responses in the UK Biobank sample (*N* = 125,925), polygenic risk scores (PRS) were generated to index 24 plausible individual risk factors for self-harm in the following domains: mental health vulnerabilities, substance use phenotypes, cognitive traits, personality traits and physical traits. PRS were entered as predictors in binomial regression models to predict self-harm. Multinomial regressions were used to model suicidal and non-suicidal self-harm. To further probe the causal nature of these relationships, two-sample Mendelian Randomisation (MR) analyses were conducted for significant risk factors identified in PRS analyses.

**Outcomes:** Self-harm was predicted by PRS indexing six individual risk factors, which are major depressive disorder (MDD), attention deficit/hyperactivity disorder (ADHD), bipolar disorder, schizophrenia, alcohol dependence disorder (ALC) and lifetime cannabis use. Effect sizes ranged from *β* = 0.044 (95% CI: 0.016 to 0.152) for PRS for lifetime cannabis use, to *β* = 0.179 (95% CI: 0.152 to 0.207) for PRS for MDD. No systematic distinctions emerged between suicidal and non-suicidal self-harm. In follow-up MR analyses, MDD, ADHD and schizophrenia emerged as plausible causal risk factors for self-harm.

**Interpretation:** Among a range of potential risk factors leading to self-harm, core predictors were found among psychiatric disorders. In addition to MDD, liabilities for schizophrenia and ADHD increased the risk for self-harm. Detection and treatment of core symptoms of these conditions, such as psychotic or impulsivity symptoms, may benefit self-harming patients.

**Funding:** Lim is funded by King’s International Postgraduate Research Scholarship. Dr Pingault is funded by grant MQ16IP16 from MQ: Transforming Mental Health. Dr Coleman is supported by the UK National Institute of Health Research Maudsley Biomedical Research Centre. MRC grant MR/N015746/1 to CML and PFO’R. Dr Hagenaars is funded by the Medical Research Council (MR/S0151132). Kylie P. Glanville is funded by the UK Medical Research Council (PhD studentship; grant MR/N015746/1). This paper represents independent research part-funded by the National Institute for Health Research (NIHR) Biomedical Research Centre at South London and Maudsley NHS Foundation Trust and King’s College London. The views expressed are those of the author(s) and not necessarily those of the NHS, the NIHR or the Department of Health and Social Care.

**Research in Context:** *Evidence before this study:* A search was conducted on PubMed for literature from inception until 1^st^ May 2019 using terms related to suicidal self-harm (SSH) and non-suicidal self-harm (NSSH), as well as polygenic risk scores (PRS), (“self-harm”[All Fields] OR “self-injurious”[All Fields] OR “self-mutilation”[All Fields] OR “suicide”[All Fields]) AND (“polygenic”[All Fields] OR “multifactorial inheritance”[All Fields]). Similar search was done for Mendelian Randomisation (MR), replacing “multifactorial inheritance” and “polygenic” with “Mendelian Randomisation/Randomization”. Evidence was included only if the study had used PRS or MR method to predict self-harm phenotypes using risk factors of self-harm. Ten papers for PRS and no paper for MR were identified. There were mixed results for PRS studies. PRS for MDD predicted SSH in two studies but not in another two studies. PRS for depressive symptoms predicted SSH but not NSSH. PRS for schizophrenia predicted SSH in one but not in another two studies. PRS for bipolar disorder predicted SSH in one study but did not predict SSH nor NSSH in another two studies.

*Added value of this study:* By using a large population-based sample, we systematically studied individual vulnerabilities and traits that can potentially lead to self-harm, including mental health vulnerabilities, substance use phenotypes, cognitive traits, personality traits and physical traits, summing up to 24 PRS as genetic proxies for 24 risk factors. We conducted MR to strengthen causal inference. We further distinguished non-suicidal self-harm (NSSH) and suicidal self-harm (SSH). Apart from PRS for schizophrenia, MDD and bipolar disorder, novel PRS were also identified to be associated with self-harm, which are PRS for attention-deficit hyperactivity disorder (ADHD), cannabis use and alcohol dependence. A larger sample size allowed us to confirm positive findings from the previously mixed literature regarding the associations between PRS for MDD, bipolar disorder, and schizophrenia with self-harm. Multivariate analyses and MR analyses strengthened the evidence implicating MDD, ADHD and schizophrenia as plausible causal risk factors for self-harm.

*Implications of all the available evidence:* Among the 24 risk factors considered, plausible causal risk factors for self-harm were identified among psychiatric conditions. Using PRS and MR methods and a number of complementary analyses provided higher confidence to infer causality and nuanced insights into the aetiology of self-harm. From a clinical perspective, detection and treatment of core symptoms of these conditions, such as psychotic or impulsivity symptoms, may prevent individuals from self-harming.

## Introduction

Self-harm is a complex trait that refers to any act of self-injury and self-poisoning carried out by an individual, regardless of intention or motivation.^1^ Being a broadly defined term, it can be further categorised into suicidal self-harm (SSH) and non-suicidal self-harm (NSSH), i.e. with or without intention of suicide. According to a meta-analysis, the cross-national prevalence rate for NSSH peaks during adolescence (17.3%), and decreases among adults (5.5%).^2^ For SSH, the cross-national prevalence rate is also the highest among adolescents (9.7%)^3^ and drops among adults (2.7%).^4^ Recently, both SSH and NSSH were included in the fifth edition of the Diagnostic and Statistical Manual of Mental Disorders (DSM-5) as separate conditions for further study.^5^ The distinction between SSH and NSSH may facilitate investigations of the aetiology and heterogeneity of self-harm.

A range of individual vulnerabilities and traits can potentially lead to self-harm, such as psychiatric illnesses,^6^ substance use,^7–9^ cognitive abilities,^10^ personality traits^11^ and physical traits.^12^ Although associations between these risk factors and self-harm have been shown in numerous observational studies, causality is difficult to infer reliably. Genetically informed designs can help in strengthening causal inference.^13^ A polygenic risk score (PRS) is a single individual-level score computed in a given trait, weighted using summary statistics from an independent genome-wide association study (GWAS) for that particular trait. A PRS for an individual risk factor (e.g. schizophrenia) can be regarded as a genetic proxy for this risk factor.^14^ To illustrate, if schizophrenia is causally related to self-harm, a PRS for schizophrenia should also be associated with self-harm. A significant association between the PRS for schizophrenia and self-harm can be regarded as an initial indication of a possible causal relationship between the two. The PRS approach can be construed as a first step in a series of genetically informed methods to investigate the aetiology of complex phenotypes, with follow-up steps including Mendelian Randomization (MR) discussed below.^14–16^

In previous studies, a PRS for major depressive disorder (MDD) was found to be associated with SSH in two clinical samples^17,18^ and one non-clinical sample.^19^ However, this was not replicated in a family-based sample.^20^ A PRS for depressive symptoms predicted SSH but not NSSH in a twin sample.^21^ On the other hand, a PRS for schizophrenia was positively associated with SSH among offspring of suicide attempters,^20^ and a population sample,^22^ but not in another clinical sample.^23^ A PRS for bipolar disorder predicted SSH in one clinical sample^24^ but did not predict SSH nor NSSH among offspring of suicide attempters,^20^ and relatives of bipolar disorder patients.^25^

The aforementioned PRS studies with mixed results were limited in several ways. Firstly, these studies focused on PRS for psychiatric disorders or psychiatric symptoms, and did not include potential risk factors from other domains, such as substance use^7–9^, cognitive abilities,^10^ personality traits^11^ and physical traits.^12^ Secondly, with two exceptions,^21,25^ none of the studies had investigated SSH and NSSH simultaneously. Thirdly, these studies have a mixture of clinical and non-clinical samples with varying sample sizes ranging from 224 individuals^23^ to 10,408 individuals,^18^ making any comparison difficult. In addition, none of these studies have implemented multivariate analyses including multiple PRS to better estimate their unique effect.

A caveat of the PRS method is its proneness to unmediated (or horizontal) pleiotropy, arising from the inclusion of many thousands of genetic variants.^14^ Unmediated pleiotropy exists when a genetic variant associated with an exposure causes the outcome through an alternative pathway, instead of via the exposure. Unmediated pleiotropy can generate associations between PRS and outcome in the absence of a causal relationship between the risk factors indexed by the PRS, and the outcome. Mendelian Randomisation (MR) can more stringently address unmediated pleiotropy and further strengthen causal inference. In MR, individual genetic variants associated with an exposure of interest are used as instrumental variables to infer causality between exposure and outcome. A number of complementary analyses, further detailed in the methods section, can be implemented to account for pleiotropy.^16^ To date, there is no published MR study which focuses on any risk factor of self-harm.

The current study will address the aforementioned limitations by systematically using 24 PRS as proxies for risk factors from different domains to predict both NSSH and SSH, using a population-based sample of 125,925 individuals. We will conduct follow-up MR analyses to strengthen causal inference.

## Methods

### Participants

The participants of the current study are a subset of the UK Biobank (http://www.ukbiobank.ac.uk). A total of 157,358 participants completed an online mental health questionnaire (MHQ) in a period from July 2016 to July 2017, which included questions regarding their lifetime symptoms of mental disorders.^26^ The participants were also genotyped. After the quality control (QC) process (see genotyping and QC details in supplementary materials), the final sample size was 125,925 individuals (56.2% females). Their ages ranged from 48 to 82 years, with a mean of 65.88 (*SD* = 7.69) years.

UK Biobank received ethical approval from the Research Ethics Committee (REC reference 11/NW/0382). The current study was conducted under the UK Biobank application 18177. Data analysis was conducted from March 2018 to June 2019.

### Defining self-harm phenotypes

To know whether the participants have ever-self-harmed, participants were asked “Have you deliberately harmed yourself, whether or not you meant to end your life?” To ascertain whether their self-harm episodes were NSSH or SSH, they were asked “Have you harmed yourself with the intention to end your life?”. In both questions, responses of “Prefer not to answer” (0.43%) were recoded as missing values. A flowchart depicting exclusion of participants and the number of participants who answered each question is shown in Figure 1.

**Figure 1.**
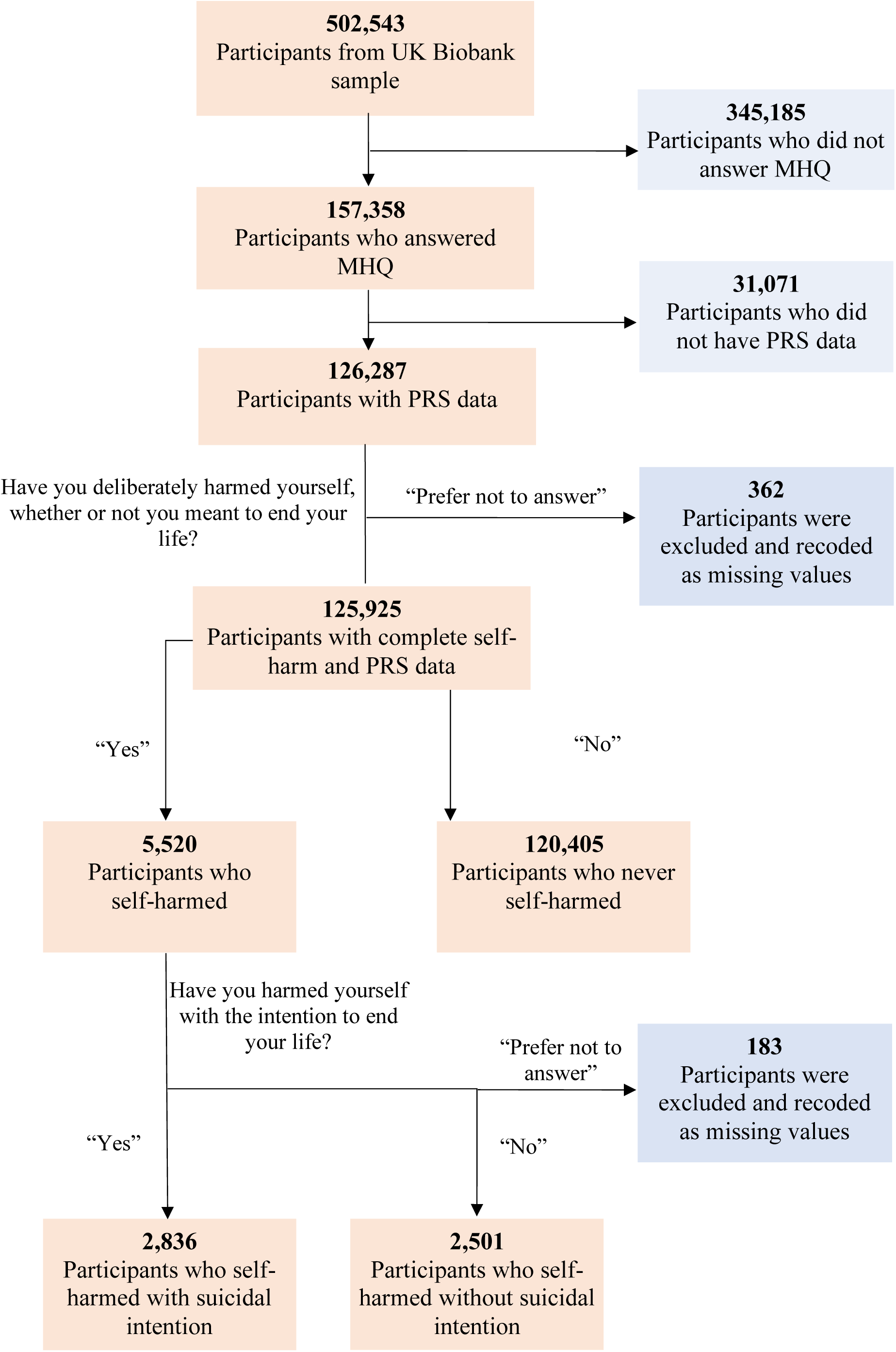
Flowchart showing the number of participants at each stage.

### Statistical Analyses

All statistical analyses were conducted in Linux environment using R version 3.5.0.^27^

### PRS analyses

PRS of UK Biobank participants were generated using PRSice-2^28^ based on their genotype data and 24 publicly available summary data from GWAS (see Table 1) selected based on the following criteria. First, we selected GWAS indexing individual vulnerabilities and traits that can potentially increase the risk of self-harm, including mental health vulnerabilities (e.g. MDD),^29^ cognitive abilities (e.g. education attainment),^30^ personality traits (e.g. neuroticism),^31^ substance use phenotypes (e.g. cannabis use),^32^ and physical traits (e.g. BMI).^12^ Second, we selected GWAS which only included participants of European ancestry and did not include UK Biobank participants (to avoid overlapping between discovery sample size and target sample). Finally, we excluded GWAS with effective sample sizes less than N = 15,000 to limit the use of underpowered PRS.

**Table 1.** Single and multiple PRS prediction of self-harm.

| Traits/disorders | Discovery<br>Sample size | Single PRS binomial model |  |  |  | Multiple PRS binomial model |  |  |
| --- | --- | --- | --- | --- | --- | --- | --- | --- |
| | | $\beta$ | 95% CI | p-value | q-value | $\beta$ | 95% CI | p-value |
| Mental health vulnerabilities |  |  |  |  |  |  |  |  |
| ADHD symptoms<sup>49</sup> | 17,666 | 0.030 | 0.003, 0.058 | 0.030 | 0.074 | - |  | - |
| ADHD<sup>50</sup> | 49,017* | 0.124 | 0.097, 0.152 | 6.69E-19 | **8.02E-18** | 0.089 | 0.062, 0.116 | **4.24E-10** |
| Alcohol dependence disorder<sup>51</sup> | 42,803* | 0.045 | 0.016, 0.073 | 2.03E-03 | **0.008** | 0.024 | -0.005, 0.053 | 0.101 |
| Anxiety disorders meta-analysis:<br>factor scores<sup>52</sup> | 18,186 | 0.026 | -0.001, 0.053 | 0.060 | 0.121 | - |  | - |
| Anxiety disorders meta-analysis:<br>case-control<sup>52</sup> | 17,310* | 0.022 | -0.005, 0.049 | 0.116 | 0.199 | - |  | - |
| Bipolar disorder<sup>53</sup> | 16,544* | 0.067 | 0.040, 0.095 | 1.74E-06 | **1.05E-05** | 0.031 | 0.002, 0.060 | **0.033** |
| MDD<sup>29</sup> | 124,331* | 0.179 | 0.152, 0.207 | 5.52E-37 | **1.33E-35** | 0.144 | 0.115, 0.173 | **3.99E-23** |
| Schizophrenia<sup>54</sup> | 75,846* | 0.128 | 0.100, 0.157 | 2.43E-18 | **1.94E-17** | 0.094 | 0.065, 0.123 | **6.99E-10** |
| Substance use phenotypes |  |  |  |  |  |  |  |  |
| Lifetime cannabis use<sup>32</sup> | 31,933* | 0.044 | 0.016, 0.072 | 0.002 | **0.008** | 0.036 | 0.009, 0.063 | **0.013** |
| Cigarettes per day<sup>55</sup> | 38,181 | -0.014 | -0.041, 0.014 | 0.329 | 0.465 | - | - | - |
| Daily alcohol use<sup>56</sup> | 70,460 | 0.001 | -0.031, 0.033 | 0.959 | 0.991 | - | - | - |
| Cognitive trait |  |  |  |  |  |  |  |  |
| Education attainment<sup>30</sup> | 106,736 | 1.69E-04 | -0.028, 0.029 | 0.991 | 0.991 | - | - | - |
| Personality traits |  |  |  |  |  |  |  |  |
| Conscientiousness<sup>57</sup> | 17,375 | -0.009 | -0.037, 0.019 | 0.524 | 0.599 | - | - | - |
| Extraversion<sup>58</sup> | 63,030 | -0.017 | -0.045, 0.010 | 0.214 | 0.343 | - | - | - |
| Neuroticism: Item Response Theory (IRT)<sup>31</sup> | 63,661 | 0·031 | 0·004, 0·058 | 0·025 | 0·074 | - | - | - |
| Agreeableness<sup>57</sup> | 17,375 | -0·011 | -0·039, 0·016 | 0·414 | 0·523 | - | - | - |
| Aggression<sup>59</sup> | 18,988 | 0·023 | -0·005, 0·050 | 0·107 | 0·198 | - | - | - |
| Antisocial behaviour<sup>60</sup> | 16,400 | 0·027 | 6·46E-05, 0·055 | 0·049 | 0·108 | - | - | - |
| **Physical traits** |  |  |  |  |  |  |  |  |
| Birth length<sup>61</sup> | 28,459 | -0·008 | -0·035, 0·020 | 0·592 | 0·646 | - | - | - |
| Birth weight<sup>62</sup> | 26,836 | 0·030 | 0·003, 0·058 | 0·031 | 0·074 | - | - | - |
| Adult height<sup>63</sup> | 253,288 | -0·018 | -0·057, 0·021 | 0·365 | 0·486 | - | - | - |
| Overweight<sup>64</sup> | 154,206* | 0·011 | -0·017, 0·038 | 0·440 | 0·527 | - | - | - |
| Extreme BMI<sup>64</sup> | 16,067* | 0·031 | 0·004, 0·059 | 0·024 | 0·074 | - | - | - |
| BMI<sup>65</sup> | 322,154 | 0·016 | -0·012, 0·044 | 0·259 | 0·388 | - | - | - |

Note: Sample size with asterisks (\*) are GWAS with case-control samples, and the effective sample sizes are calculated using the formula $N_{\text{effective}} = 4 / (1/N_{\text{cases}} + 1/N_{\text{controls}})$ whenever possible. The $p$ -values or $q$ -values in bold are those that met the nominal $p < .05$ or corrected $q < .05$ thresholds.

Each participant had 24 PRS, which were each calculated as the sum of alleles associated with their respective phenotypes, weighted by their effect sizes with *p*-values less than a threshold *p*_*T*_ < 0.3 (selecting and reporting results from a single threshold allowed us to limit multiple testing, as done in previous PRS studies).^22,33^ Clumping was used to remove SNPs in linkage equilibrium (*r*^*2*^ < 0.1 within a 250 kb window). All PRS in the final analytical sample were standardised.

#### Single PRS Binomial Logistic Regression

For each PRS, a binomial logistic regression was conducted to test whether it predicted self-harm (i.e. “Self-harmed” versus “Never self-harmed”).

#### Multiple PRS Binomial Logistic Regression

All PRS significantly associated with self-harm in single PRS binomial logistic regressions were then jointly modelled in a multivariate binomial logistic regression model to assess their unique effects.

#### Multinomial Logistic Regressions

To investigate whether each PRS differentially predicted NSSH versus SSH, we fitted a series of multinomial logistic regression models. We first compared each of the NSSH and SSH groups to the never self-harmed group (i.e. “Never self-harmed” as the reference group). We then directly compared NSSH and SSH by testing a model with “NSSH” as the reference group.

#### Covariates and multiple testing

All regression models were controlled for sex, age and population stratification (by including assessment centre, genotyping batch and the first 6 principal components as covariates in the models). To control for multiple testing in single PRS binomial and multinomial regressions, we employed the false discovery rate (FDR) method^34^ which controls the expected proportion of false positives among the rejected hypotheses. We used *q* < .05 as the significance threshold.

### MR analyses

All MR analyses were conducted using R package TwoSampleMR.^35^ Risk factors for which their PRS significantly predicted self-harm were selected for follow-up MR analyses. For self-harm in UK Biobank sample as the outcome for MR analyses, we obtained GWAS summary statistics from Neale Lab (http://www.nealelab.is/uk-biobank). SNPs of the exposures which passed the p-value threshold of *p* < 5E-5 were selected as instrumental variables for MR analyses. A liberal threshold was used to ensure that enough variants were available for all risk factors, including those with few genome-wide significant SNPs (e.g. ADHD). The strategy entails potential weak instrument bias. In two-sample MR, the resulting bias is towards the null, making estimates more conservative (see below how this was dealt with).^36^ Clumping of SNPs with r^2^ < .001 within 250 kb was applied. SNPs in exposures and outcomes were harmonized by flipping alleles where possible, and we use allele frequencies to infer strands of ambiguous SNPs. Non-inferable SNPs with minor allele frequency > 0.42 were discarded.

We selected four MR methods which have different strengths and limitations. We conducted univariable MR using:

i. Inverse variance weighted (IVW) method, which is the most powerful method but cannot account for directional pleiotropy;^37^
ii. Robust Adjusted Profile Score (RAPS) method, which is used to account for the selection of weak instruments;^33^
iii. Weighted median method, as it is more robust to directional pleiotropy than IVW and is more robust to individual genetic variants with outlying causal estimates than IVW and MR-Egger;^38^ and
iv. MR-Egger regression method, whereby significance of its intercept term can inform on the presence of directional pleiotropy.^39^

In addition, MR Steiger filtering^40^ was implemented to address the possibility of reverse causation (i.e. self-harm causing the putative risk factor). For each SNP, we expect that the effect size for the association with the exposure should be larger than the effect size for the association with the outcome. This is because the effect on the outcome is hypothesised to be indirect through the exposure. As such, all SNPs for which the effect size of the association with the outcome was larger than the one with the exposure were filtered out before reimplementing MR. Finally, similar to PRS analyses, exposures which were significant in univariable MR were assessed for their independent effect in a multivariable MR model using the IVW method.

For PRS analyses, we conducted further complementary analyses excluding cases with MDD and schizophrenia diagnoses to investigate the effect of genetic liability on self-harm with the influence of diagnoses excluded. We also calculated risk ratios for medicated and non-medicated cases compared to those with median PRS in the general population (see supplementary materials for definitions of cases and medication). We created a quantile plot separating the participants into three groups: general population (in 20 quantiles), medicated cases and unmedicated cases, and calculated the risk ratios of these groups for self-harm relative to the group in the population with median PRS.

## Results

### Descriptive statistics

Figure 1 shows the number of participants who: never self-harmed, self-harmed, engaged in SSH, and engaged in NSSH. Table 2 shows the gender proportion, and mean age of each subgroup.

**Table 2.** Descriptive statistics for each subgroup of self-harm related phenotypes.

| Subgroup of sample | Female (%) | Mean age (years) | SD of age (years) |
| --- | --- | --- | --- |
| Full analytical sample | 56·2 | 65·9 | 7·7 |
| Self-harmed | 69·4 | 62·3 | 7·5 |
| SSH | 68·0 | 63·1 | 7·4 |
| NSSH | 70·5 | 61·4 | 7·5 |
| Never self-harmed | 55·6 | 66·1 | 7·7 |

### PRS analyses

#### Single PRS binomial logistic regression

Table 1 and Figure 2 show results from 24 single PRS binomial logistic regression tests, using each PRS as a predictor variable. Out of the 24 PRS, 10 PRS were significant predictors of self-harm at the nominal level (p < 0.05). After applying FDR correction, 6 PRS had *q*-value < .05. In order of decreasing effect sizes, they are PRS for: MDD, schizophrenia, ADHD, bipolar disorder, alcohol dependence disorder (ALC), and lifetime cannabis use, with effect sizes ranging from *β* = 0.179 (95% CI: 0.152 to 0.207) for MDD, to *β* = 0.044 (95% CI: 0.016 to 0.072) for lifetime cannabis use. Figure S3 shows the pseudo R^2^ plots of these 6 PRS in accounting for the variance in self-harm.

**Figure 2.**
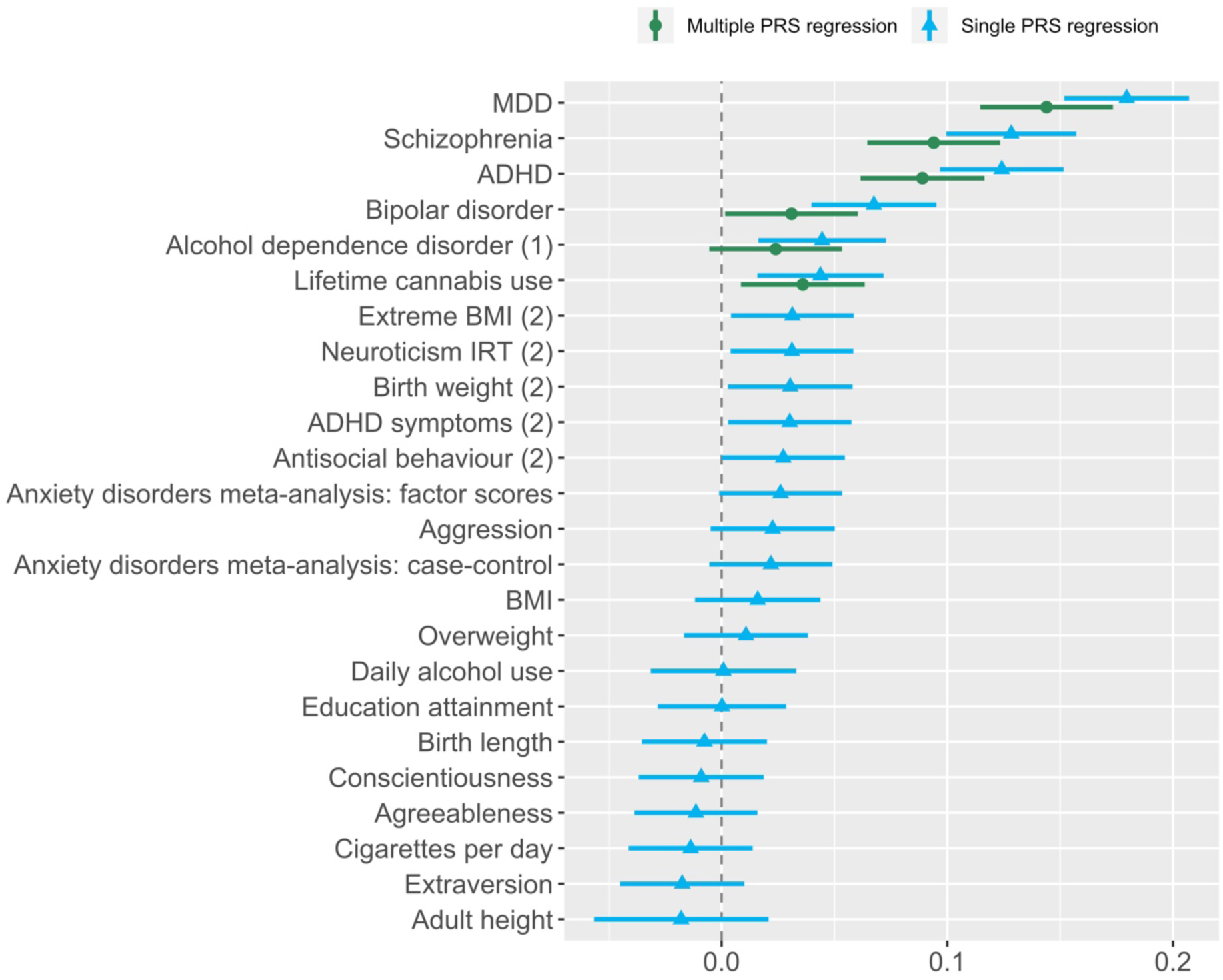
Estimates from single PRS regression and multiple PRS regression in decreasing effect sizes. (1) indicates not significant in multiple PRS regression. (2) indicates not significant after FDR correction.

#### Multiple PRS binomial logistic regression

In the multiple PRS model, all PRS except the PRS for ALC had an independent effect of self-harm as shown in Table 1 and Figure 2. By controlling for the effects of other PRS, effect sizes of these PRS have diminished slightly compared to those in single PRS binomial logistic regression, ranging from *β* = 0.144 (95% CI: 0.115 to 0.173) for MDD to *β* = 0.031 (95% CI: 0.002 to 0.060) for bipolar disorder. These PRS were weakly correlated, ranging from *r* = 0.01 (between bipolar disorder and ADHD) to *r* = 0.22 (between schizophrenia and bipolar disorder; see Table S3 for all correlations), suggesting that multicollinearity was not an issue.

#### Single PRS multinomial logistic regression

Table S1 shows results from 24 multinomial logistic regression tests, using PRS as predictor variable for three possible outcomes: “Never self-harmed”, “NSSH” and “SSH”. When “Never self-harmed” was used as the reference group, PRS for bipolar disorder, lifetime cannabis use and extreme BMI predicted SSH but not NSSH, with *q* < .05. However, when “NSSH” was set as the reference group in order to directly compare NSSH versus SSH, none of the PRS significantly distinguished between NSSH versus SSH.

### MR analyses

Table 3 shows the results from MR analyses. ADHD, ALC, bipolar disorder, lifetime cannabis use, MDD and schizophrenia were exposures in 6 separate univariable MR analyses, with self-harm as the outcome. Out of these 6 exposures, MDD, ADHD and schizophrenia had MR estimates with *p*-values < .05. For other exposures, none of their MR estimates had *p* < .05.

**Table 3.** Univariable and multivariable MR analyses

| Exposure | Univariable MR |  |  |  |  |  | Multivariable MR |  |  |  |  |
| --- | --- | --- | --- | --- | --- | --- | --- | --- | --- | --- | --- |
|  | method | nsnp | b | LowCI | UpCI | pval | nsnp | b | lowCI | UpCI | p-value |
| ADHD | IVW | 244 | 0.003 | 0.001 | 0.005 | **0.001** | 1206 | 0.001 | -0.001 | 0.003 | 0.269 |
|  | MR RAPS |  | 0.003 | 0.001 | 0.005 | **0.001** |  |  |  |  |  |
|  | Weighted median |  | 0.003 | 2.83E-04 | 0.005 | **0.028** |  |  |  |  |  |
|  | MR Egger |  | 0.006 | 0.001 | 0.011 | **0.031** |  |  |  |  |  |
|  | MR Egger intercept |  | -2.57E-04 | -6.88E-04 | 1.74E-04 | 0.243 |  |  |  |  |  |
| Alcohol dependence disorder | IVW | 86 | 0.001 | -4.50E-04 | 0.003 | 0.157 | - | - | - | - | - |
|  | MR RAPS |  | 0.001 | -4.57E-04 | 0.003 | 0.150 |  |  |  |  |  |
|  | Weighted median |  | 0.001 | -0.001 | 0.004 | 0.332 |  |  |  |  |  |
|  | MR Egger |  | 0.001 | -0.002 | 0.005 | 0.465 |  |  |  |  |  |
|  | MR Egger intercept |  | -1.88E-05 | -5.85E-04 | 5.48E-04 | 0.948 |  |  |  |  |  |
| Bipolar disorder | IVW | 77 | 0.001 | -0.001 | 0.003 | 0.310 | - | - | - | - | - |
|  | MR RAPS |  | 0.001 | -0.001 | 0.003 | 0.355 |  |  |  |  |  |
|  | Weighted median |  | 1.27E-04 | -0.002 | 0.003 | 0.922 |  |  |  |  |  |
|  | MR Egger |  | 3.22E-04 | -0.008 | 0.008 | 0.936 |  |  |  |  |  |
|  | MR Egger intercept |  | 8.31E-05 | -9.39E-04 | 1.11E-03 | 0.874 |  |  |  |  |  |
| Lifetime cannabis use | IVW | 85 | -2.07E-04 | -0.002 | 0.001 | 0.787 | - | - | - | - | - |
|  | MR RAPS |  | -2.85E-04 | -0.002 | 0.001 | 0.730 |  |  |  |  |  |
|  | Weighted median |  | -0.001 | -0.004 | 0.001 | 0.314 |  |  |  |  |  |
|  | MR Egger |  | -0.002 | -0.006 | 0.001 | 0.171 |  |  |  |  |  |
|  | MR Egger intercept |  | 3.62E-04 | -1.38E-04 | 8.61E-04 | 0.160 |  |  |  |  |  |
| MDD | IVW | 239 | 0.008 | 0.005 | 0.011 | **2.84E-08** | 1206 | 0.011 | 0.007 | 0.015 | **1.04E-12** |
|  | MR RAPS |  | 0.008 | 0.005 | 0.011 | **1.24E-07** |  |  |  |  |  |
|  | Weighted median |  | 0.006 | 0.001 | 0.011 | **0.013** |  |  |  |  |  |
|  | MR Egger |  | 0.002 | -0.004 | 0.008 | 0.463 |  |  |  |  |  |
|  | MR Egger intercept |  | 4.20E-04 | 6.06E-05 | 7.80E-04 | **0.023** |  |  |  |  |  |
| Schizophrenia | IVW | 1003 | 0.003 | 0.002 | 0.004 | **1.54E-09** | 1206 | 0.002 | 4.00E-05 | 0.004 | **0.002** |
|  | MR RAPS |  | 0.003 | 0.002 | 0.004 | **2.89E-09** |  |  |  |  |  |
|  | Weighted median |  | 0.003 | 0.002 | 0.005 | **1.80E-05** |  |  |  |  |  |
|  | MR Egger |  | 0.004 | 0.001 | 0.007 | **0.009** |  |  |  |  |  |
|  | MR Egger intercept |  | -4.84E-05 | -2.42E-04 | 1.45E-04 | 0.625 |  |  |  |  |  |

Note. The $p$ -values in bold are those that met the $p < .05$ threshold.

For MDD, despite having the strongest IVW (*β* =0.008, 95% CI: 0.005 to 0.011, *p* = 2.84E-08), MR RAPS (*β* =0.008, 95% CI: 0.005 to 0.011, *p* = 1.24E-07), and weighted median (*β* = 0.006, 95% CI: 0.001 to 0.011, *p* = 0.013) estimates among the three exposures, the MR Egger estimate was not significant. On the other hand, all MR estimates for ADHD and schizophrenia were significant.

The significance of intercept terms in MR-Egger test indicates the presence of pleiotropy. Out of the 6 exposures in MR-Egger test, none of the intercept terms were significant, except for MDD (*p* = 0.023). MR Steiger directionality tests could only be applied to test the direction of causality between ADHD, MDD and schizophrenia with self-harm because the summary statistics for other exposures did not contain information about allele frequencies, which are needed for the test. MR Steiger directionality test showed that all SNPs of MDD, schizophrenia and ADHD are more predictive of the respective exposures than self-harm, suggesting that reverse causation unlikely explained our findings.

When ADHD, MDD and schizophrenia were included as exposures in multivariable IVW MR analysis, only MDD (*β* = 0.011, 95% CI: 0.007 to 0.015, *p* = 1.04E-12) and schizophrenia (*β* = 0.002, 95% CI: 4.00E-05 to 0.004, *p* = 0.002) remained as independent predictors of self-harm. Due to the potential presence of pleiotropy between MDD and self-harm, another multivariable IVW MR model was conducted with only ADHD and schizophrenia as exposures. Both ADHD (*β* = 0.003, 95% CI: 0.001 to 0.005, *p* = 2.21E-04) and schizophrenia (*β* = 0.003, 95% CI: 0.002 to 0.004, *p* = 7.60E-07) were significant predictors in this model.

In PRS complementary analyses which excluded cases, PRS for MDD and schizophrenia still predicted self-harm in a healthy, screened cohort, indicating that genetic liabilities can predict self-harm when influence of diagnoses is excluded (See Table S2). In the quantile plot, cases for schizophrenia and MDD appear to be at much larger risk for self-harm than the rest of the population. Medicated MDD cases were at higher risk of self-harm than non-medicated MDD cases, which was not the case for schizophrenia (See Figure 3).

**Figure 3.**
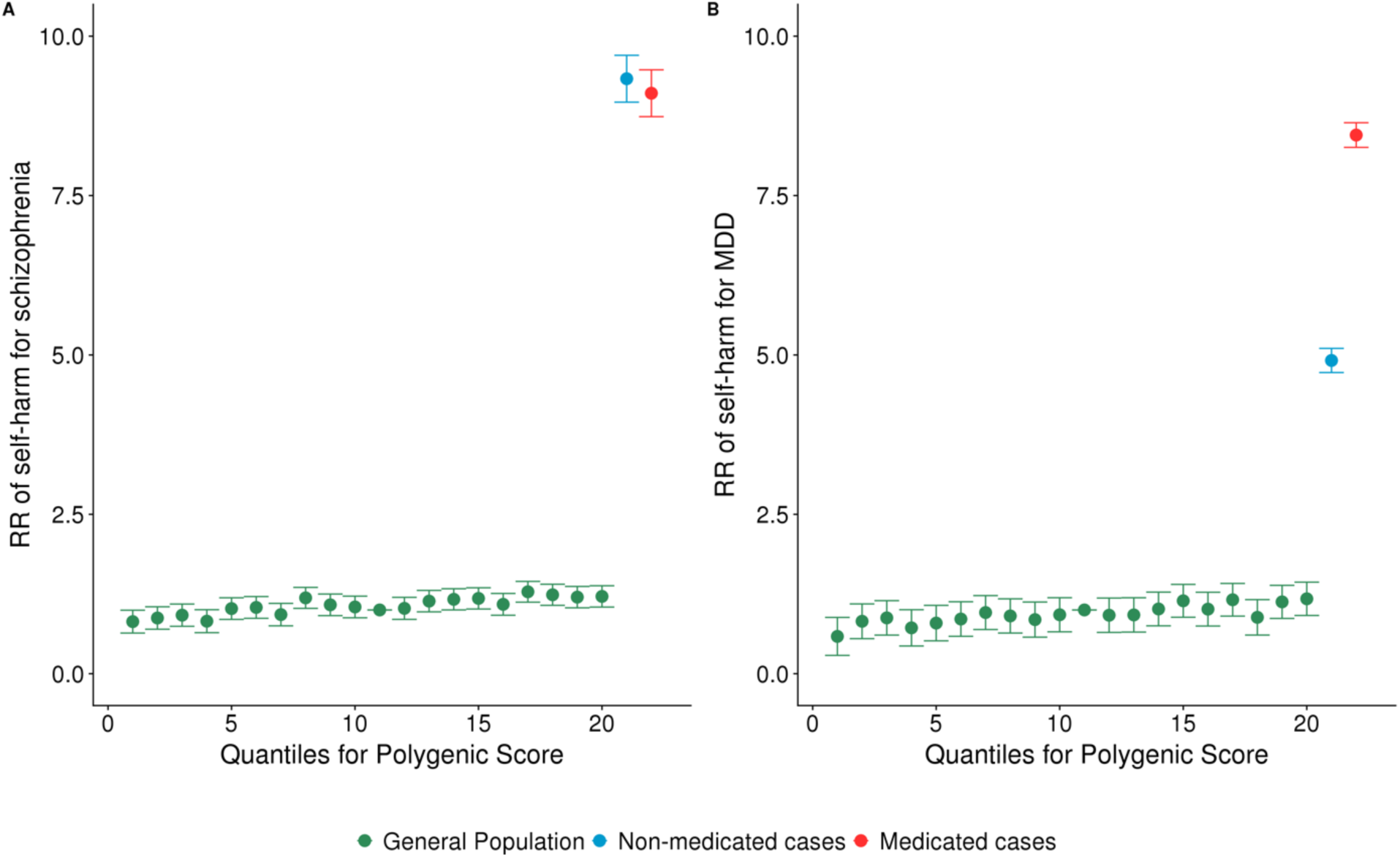
Relative risks of general population, non-medicated cases and medicated cases in self-harming compared to those with median PRS (11^th^ quantile) in schizophrenia (A) and MDD (B). Out of 177 schizophrenia cases in the final analytical sample, 89 (50.3%) of them were medicated. Out of 34,680 MDD cases in the final analytical sample, 7,852 (22.6%) of them were medicated.

## Discussion

To our knowledge, this is the first study using multiple PRS as genetic proxies to systematically investigate a range of individual vulnerabilities and traits as risk factors for self-harm in a large population sample. In PRS analyses, we identified 6 risk factors (i.e. MDD, schizophrenia, ADHD, bipolar disorder, ALC, and lifetime cannabis use) which predicted self-harm. Five among six (except for ALC) remained significant in a multiple PRS regression. We found no evidence of differential prediction for SSH versus NSSH. In follow-up MR analyses, MDD, schizophrenia and ADHD emerged as plausible causal risk factors for self-harm, despite evidence of unmediated pleiotropy for MDD. We discuss in turn: (1) insights into the aetiology of self-harm, and (2) clinical implications.

### Insights into the aetiology of self-harm

Results from our PRS methods corroborated previous observational findings where MDD,^6^ schizophrenia,^41^ ADHD,^42^ bipolar disorder,^6^ and ALC^8^ were phenotypically associated with self-harm. Our results are also consistent with positive associations found in PRS studies for MDD,^17,19,43^ schizophrenia,^20^ and bipolar disorder^24^. Previous mixed findings for these PRS may have stemmed from lack of power, as sample sizes for those studies varied widely. The current study adds lifetime cannabis use, ADHD, and ALC as novel PRS associated with self-harm. However, when controlling for other PRS, the PRS for ALC did not significantly predict self-harm. This finding may suggest that the genetic liability for ALC does not independently predict self-harm when the effect of genetic liability for MDD, bipolar disorder, schizophrenia, ADHD and lifetime cannabis are accounted for. For example, ALC may be a marker for a true predictor such as impulsivity which is more efficiently captured in the PRS for ADHD.^44^. Alternatively, null findings for ALC can also plausibly be due to a lack of power compared to other polygenic scores. Hence, we cannot completely rule out that the PRS for ALC has an independent effect on self-harm and the corresponding causal effect of ALC on self-harm.

Most of the PRS which predicted self-harm in the current study relate to psychiatric conditions, which confirms the prominence of psychiatric conditions in the aetiology of self-harm.^45^ Beyond psychiatric conditions, cognitive traits, physical traits, and personality traits were not found to be associated with self-harm using PRS approach, although previous observational findings found significant phenotypic associations for these three domains.^10–12^ The absence of significant findings in this case is unlikely to be solely due to lack of power, given that GWAS for some of these traits are more powerful than GWAS for psychiatric conditions (e.g. BMI and education attainment). These findings suggest that these traits and vulnerabilities are unlikely to have (strong) causal effects on self-harm.

Our MR analyses provided further support for the role of MDD, ADHD, and schizophrenia in the aetiology of self-harm. An intriguing finding is the presence of significant pleiotropy in the case of MDD. Rather than signifying that MDD does not have a causal effect on self-harm, this may reflect a possible measurement issue. Indeed, one of the diagnostic criteria for MDD is related to having suicidal thoughts and attempts, which could artificially introduce a pleiotropic effect.^5^ To deal with this issue, future studies may rely on a GWAS for MDD excluding the diagnostic criteria related to suicidal thoughts and attempts. This might also explain why, in multivariate MR, the effect of ADHD was no longer significant – as we partially controlled for self-harm – whereas it was significant when only considering ADHD and schizophrenia.

The current study found mixed results for whether there are distinct aetiologies for SSH and NSSH. Most PRS which predicted self-harm also predicted both SSH and NSSH, except bipolar disorder, lifetime cannabis use and extreme BMI, which only predicted SSH but not NSSH from those who never self-harmed. However, in a formal test comparing NSSH and SSH, the estimates of these three risk factors were not significantly different between NSSH and SSH. Hence, our findings do not provide evidence for marked differences in aetiology between SSH and NSSH.

### Clinical implications

The current study suggests that individual vulnerabilities and traits underlying self-harm most likely relate to psychiatric conditions such as MDD and schizophrenia, rather than to other domains such as personality traits. Hence, treatments focusing on the core symptoms of these psychiatric conditions are important in preventing or addressing the risk of self-harm. Findings from PRS analyses suggest that genetic liabilities for these conditions increase the likelihood of self-harm even in those not clinically diagnosed. This may suggest that subthreshold symptoms of these core psychiatric conditions may increase the risk of self-harm. Clinicians may want to systematically test for such symptoms in self-harming patients. Future investigations may test whether drugs for such core conditions may be repurposed for treating self-harming patients, with either full blown or subthreshold conditions. For example, prescription of methylphenidate for ADHD treatment was found to be associated with reduction of suicide attempt risk.^46^ As a note of caution, treated schizophrenia cases were not at less risk of self-harm than non-treated patients whereas treated MDD patients were at substantial higher risk for self-harm. This could be due to treated patients having more severe symptoms than untreated patients, or it could be due to adverse effects of medication, in particular in the case of MDD where suicidality might be an adverse effect of antidepressant treatment.^47^

### Limitations

In order to avoid the overlapping of discovery and target sample, we excluded GWAS which contain UK Biobank sample, resulting in selecting older GWAS for generating PRS in some cases. This might have led to non-significant findings due to lack of power. The results should be generalised with caution because UK Biobank is not representative of the UK population as they are more educated, older, wealthier, and healthier.^48^ The questions asked in MHQ were retrospective and their formulation led to an exclusive dichotomy between NSSH or SSH, whereby some might have engaged in both NSSH and SSH at different times.

### Conclusion

Among 24 PRS used as genetic proxies for vulnerabilities and traits possibly associated with self-harm, we found that PRS for MDD, schizophrenia, ADHD, bipolar disorder, ALC and cannabis were statistically significant. After a series of complementary analyses to further strengthen the causal inference, schizophrenia survived as the most plausible causal risk factor, followed by MDD and ADHD. Detection and treatment of core symptoms of these conditions, such as psychotic or impulsivity symptoms, may benefit self-harming patients.

