## Supplementary materials for "Identifying Potential Causal Risk Factors for Self-Harm: A Polygenic Risk Scoring and Mendelian Randomisation Approach"

**Genotyping, imputation and quality control**

Participants were genotyped across 22 assessment centres in the UK, using either the Affymetrix UK BiLEVE Axiom array or the Affymetrix UK Biobank Axiom® array.[1] These two arrays share over 95% common content. Further details of genotyping and sample processing are available (http://biobank.ctsu.ox.ac.uk/crystal/docs/genotyping_affy_sampro.pdf). Before the release of UK Biobank genetic data, a stringent quality control (QC) protocol was performed, with further imputation[1] using the Haplotype Reference Consortium (HRC) reference panel.[2] Details of the QC protocol (http://biobank.ctsu.ox.ac.uk/crystal/docs/genotyping_qc.pdf) and genetic imputation (http://www.ukbiobank.ac.uk/wp-content/uploads/2014/04/imputation_documentation_May2015.pdf) are available online.

Prior to the analyses in the current study, further QC steps were taken. Participants were excluded if they were reported as outliers based on heterozygosity and missing rates, had a variant call rate < 98%, had aa mismatch of phenotypic and genotypic gender, or had non-European ancestry as defined by 4-means clustering of the first two PCs from the genetic data. By using the KING toolset,[3] one of each pair of participants sharing relatedness of up to the third degree (KING estimated kinship coefficient > 0.044) were also excluded. Removal of relatives was performed using a "greedy" algorithm, which minimises exclusions (for example, by excluding the child in a mother-father-child trio). Genetic variants were excluded if they had a call rate < 98%, a minor allele frequency < 0.01, or deviated from the Hardy-Weinberg equilibrium ($p < 10^{-8}$). Principal components analysis was also performed on the European-only subset of the data using the software flashpca2.[4]

### Complementary analyses

*Definition of cases*

Cases for MDD and schizophrenia were identified respectively from UK Biobank's primary and secondary ICD 9 or ICD 10 diagnoses, non-cancer illness self-report and MHQ self-report. The details are shown in the table below. Number of cases derived from each data field are presented in Figures S3 (for MDD) and S4 (for schizophrenia).

| Phenotype | Data field |
|---|---|
| ICD 9 diagnosis (primary) | 41203 |
| ICD 9 diagnosis (secondary) | 41205 |
| ICD 10 diagnosis (primary) | 41202 |
| ICD 10 diagnosis (secondary) | 41204 |
| Non-cancer illness self-report | 20002 |
| MHQ self-report ("Have you been diagnosed with one or more of the following mental health problems by a professional, even if you don't have it currently?") | 20544 |

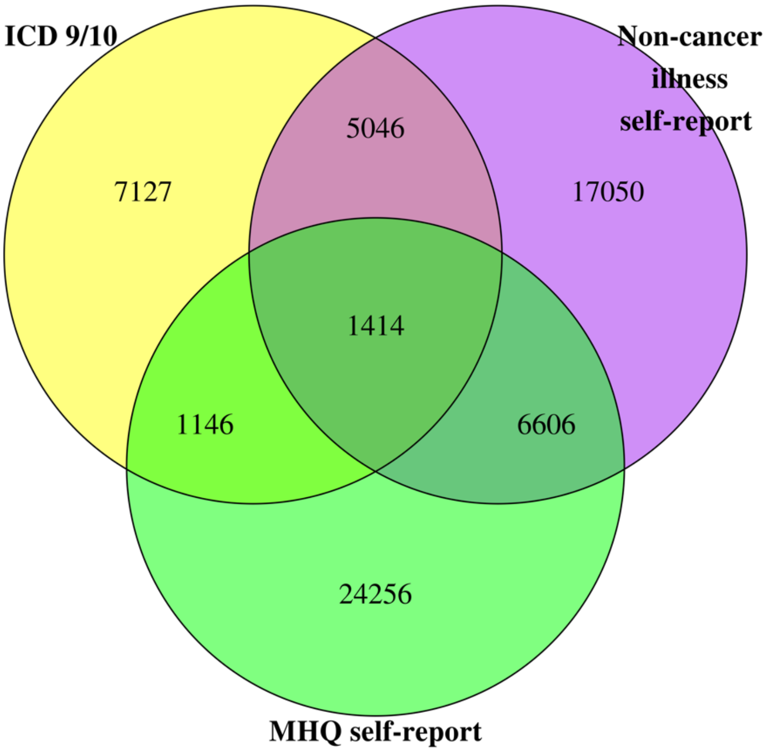

Figure S1. Number of MDD cases derived from respective data fields.

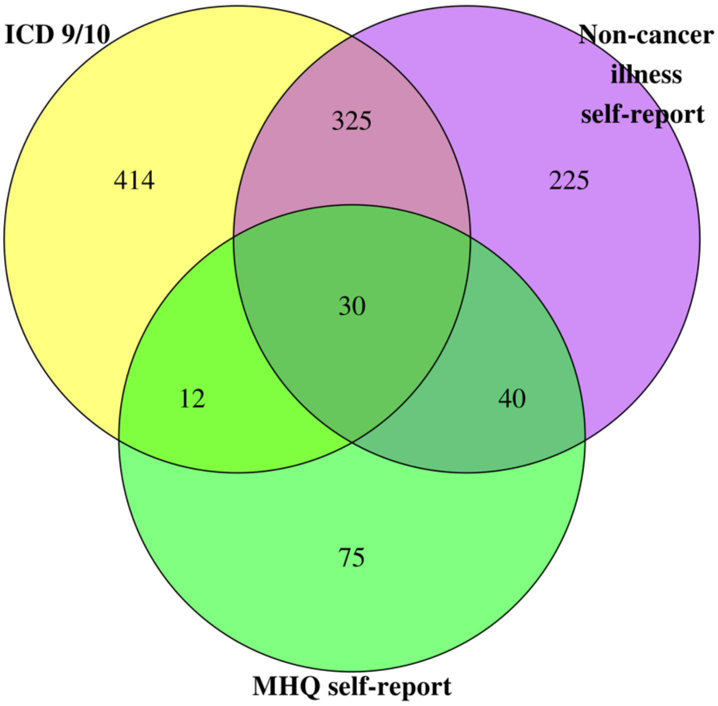

Figure S2. Number of schizophrenia cases derived from respective data fields.

Two complementary analyses were conducted for risk factors which were significant exposures for self-harm in multivariate MR analyses. Firstly, single PRS binomial regression tests were repeated for these risk factors by excluding participants who are associated with these risk factors (i.e. having diagnoses for MDD or schizophrenia). This was to test if the association between their respective PRS and self-harm remains after those with the respective psychiatric disorder diagnoses were excluded.

*Definition of medication*

Lists of antidepressants and antipsychotics prescribed for MDD and schizophrenia patients were compiled from the Mind website (for antidepressants: https://www.mind.org.uk/information-support/drugs-and-treatments/antidepressants-a-z/#.XP-iKy3Mw1I; for antipsychotics: https://www.mind.org.uk/information-support/drugs-and-treatments/antipsychotics-a-z/overview/?o=60249#.XP-iUS3Mw1I ). We also visited British National Formulary (BNF) website (https://www.bnf.org) to identify other medications used for MDD or schizophrenia. Data of medications taken by the participants were derived from UK Biobank's treatment/medication code (data field 20003).

### Results

Table S1. Single PRS binomial regressions with cases excluded.

| PRS (with cases removed) | *ß* | 95% CI | *p*-value |
|---|---|---|---|
| Schizophrenia | 0·123 | 0·094, 0·152 | 6·78E-17 |
| MDD | 0·138 | 0·093, 0·183 | 1·96E-09 |

Table S2. Multinomial regression models

| PRS of risk factors | NSSH vs never self-harmed[a] | | | | SSH vs never self-harmed[b] | | | | NSSH vs SSH[c] | | | |
|---|---|---|---|---|---|---|---|---|---|---|---|---|
| | *ß* | 95% CI | p-value | q-value | *ß* | 95% CI | p-value | q-value | *ß* | 95%CI | p-value | q-value |
| **Mental Health Vulnerabilities** | | | | | | | | | | | | |
| ADHD symptoms | 0·042 | 0.002, 0.082 | 0·041 | 0·396 | 0·034 | -0.003, 0.072 | 0·073 | 0·250 | 0·004 | -0.050, 0.058 | 0·881 | 1·000 |
| ADHD | 0·125 | 0.085, 0.165 | 9·98E-10 | **2·4E-08** | 0·113 | 0.075, 0.150 | 3·96E-09 | **6·34E-08** | -0·008 | -0.062, 0.046 | 0·767 | 1·000 |
| Alcohol dependence disorder | 0·036 | -0.006, 0.077 | 0·091 | 0·548 | 0·048 | 0.009, 0.087 | 0·015 | 0·083 | -0·007 | -0.062, 0.048 | 0·801 | 1·000 |
| Anxiety disorders meta-analysis: factor scores | 0·043 | 0.002, 0.083 | 0·038 | 0·396 | 4·32E-04 | -0.037, 0.038 | 0·982 | 1·000 | -0·048 | -0.102, 0.006 | 0·079 | 0·597 |
| Anxiety disorders meta-analysis: case-control | -0·001 | -0.041, 0.039 | 0·948 | 1·000 | 0·043 | 0.005, 0.080 | 0·026 | 0·122 | 0·042 | -0.012, 0.095 | 0·131 | 0·656 |
| Bipolar disorder | 0·036 | -0.005, 0.076 | 0·086 | 0·548 | 0·099 | 0.061, 0.137 | 3·20E-07 | **3·84E-06** | 0·055 | 0.001, 0.109 | 0·045 | 0·546 |
| MDD | 0·157 | 0.116, 0.197 | 3·33E-14 | **1·60E-12** | 0·198 | 0.159, 0.236 | 2·20E-16 | **2·20E-16** | 0·046 | -0.008, 0.101 | 0·094 | 0·597 |
| Schizophrenia | 0·123 | 0.081, 0.165 | 8.64E-09 | **1·38E-07** | 0·133 | 0.094, 0.173 | 3·27E-11 | **7·84E-10** | 0·009 | -0.046, 0.064 | 0·756 | 1·000 |
| **Substance use phenotypes** | | | | | | | | | | | | |
| Lifetime cannabis use | 0·033 | -0.008, 0.074 | 0·110 | 0·586 | 0·056 | 0.017, 0.094 | 0·005 | **0·036** | 0·014 | -0.040, 0.069 | 0·612 | 1·000 |
| Cigarettes per day | -0·025 | -0.064, 0.015 | 0·229 | 0·845 | -0·009 | -0.047, 0.029 | 0·643 | 1·000 | 0·010 | -0.045, 0.064 | 0·732 | 1·000 |
| Daily alcohol use | -0·005 | -0.052, 0.042 | 0·827 | 1·000 | 0·003 | -0.041, 0.048 | 0·883 | 1·000 | 0·006 | -0.056, 0.069 | 0·843 | 1·000 |

| | | | | | | | | | | | | |
|---|---|---|---|---|---|---|---|---|---|---|---|---|
| **Cognitive trait** | | | | | | | | | | | | |
| Education attainment | 0·040 | -0.003, 0.082 | 0·068 | 0·547 | -0·014 | -0.053, 0.025 | 0·478 | 1·000 | -0·088 | -0.143, -0.033 | 0·002 | 0·086 |
| **Personality traits** | | | | | | | | | | | | |
| Conscientiousness | -0·002 | -0.043, 0.039 | 0·915 | 1·000 | -0·025 | -0.063, 0.013 | 0·194 | 0·583 | -0·022 | -0.076, 0.033 | 0·438 | 1·000 |
| Extraversion | 0·011 | -0.030, 0.051 | 0·601 | 1·000 | -0·033 | -0.071, 0.005 | 0·089 | 0·284 | -0·041 | -0.095, 0.013 | 0·137 | 0·656 |
| Neuroticism IRT | 0·030 | -0.009, 0.070 | 0·133 | 0·638 | 0·046 | 0.009, 0.084 | 0·016 | 0·083 | 0·022 | -0.032, 0.076 | 0·429 | 1·000 |
| Agreeableness | -0·022 | -0.062, 0.018 | 0·287 | 0·983 | -0·009 | -0.046, 0.029 | 0·656 | 1·000 | -0·002 | -0.056, 0.052 | 0·950 | 1·000 |
| Aggression | 0·026 | -0.015, 0.066 | 0·211 | 0·845 | 0·013 | -0.024, 0.051 | 0·487 | 1·000 | 0·011 | -0.043, 0.066 | 0·685 | 1·000 |
| Antisocial behaviour | 0·015 | -0.025, 0.054 | 0·462 | 1·000 | 0·040 | 0.003, 0.078 | 0·036 | 0·143 | 0·011 | -0.043, 0.065 | 0·699 | 1·000 |
| **Physical traits** | | | | | | | | | | | | |
| Birth length | 0·025 | -0.016, 0.066 | 0·227 | 0·845 | -0·042 | -0.080, -0.005 | 0·028 | 0·122 | -0·063 | -0.117, -0.008 | 0·024 | 0·391 |
| Birth weight | 0·020 | -0.021, 0.060 | 0·344 | 1·000 | 0·037 | -3.34E-04, 0.075 | 0·052 | 0·192 | 0·019 | -0.036, 0.074 | 0·498 | 1·000 |
| Adult height | -0·028 | -0.085, 0.029 | 0·332 | 1·000 | -0·032 | -0.085, 0.020 | 0·230 | 0·649 | -0·043 | -0.095, 0.008 | 0·099 | 0·597 |
| Overweight | -0·005 | -0.045, 0.036 | 0·823 | 1·000 | 0·017 | -0.020, 0.055 | 0·368 | 0·982 | 0·019 | -0.035, 0.073 | 0·498 | 1·000 |
| Extreme BMI | 0·006 | -0.034, 0.046 | 0·761 | 1·000 | 0·055 | 0.017, 0.092 | 0·004 | **0·036** | 0·050 | -0.004, 0.104 | 0·067 | 0·597 |
| BMI | -0·018 | -0.058, 0.023 | 0·398 | 1·000 | 0·051 | 0.013, 0.089 | 0·009 | 0·062 | 0·072 | 0.018, 0.127 | 0·009 | 0·228 |

*Note.* [a,b] These are the outputs from the same multinomial model, in which “Never self-harmed” was the reference group. [c] These outputs were derived from a different multinomial model, in which “NSSH” was the reference group. The *q*-values in bold are those that met the $q < .05$ threshold.

Table S3. Correlations between PRS which were significant in single PRS regression.

| | ADHD | ADD | Bipolar disorder | Lifetime Cannabis use | MDD | Schizophrenia |
|---|---|---|---|---|---|---|
| ADHD | 1.00 | 0·05 | 0·01 | 0·02 | 0·21 | 0·06 |
| ADD | 0·05 | 1.00 | 0·05 | 0·07 | 0·06 | 0·08 |
| Bipolar disorder | 0·01 | 0·05 | 1.00 | 0·04 | 0·10 | 0·22 |
| Lifetime Cannabis use | 0·02 | 0·07 | 0·04 | 1.00 | 0·04 | 0·08 |
| MDD | 0·21 | 0·06 | 0·10 | 0·04 | 1.00 | 0·16 |
| Schizophrenia | 0·06 | 0·08 | 0·22 | 0·08 | 0·16 | 1.00 |

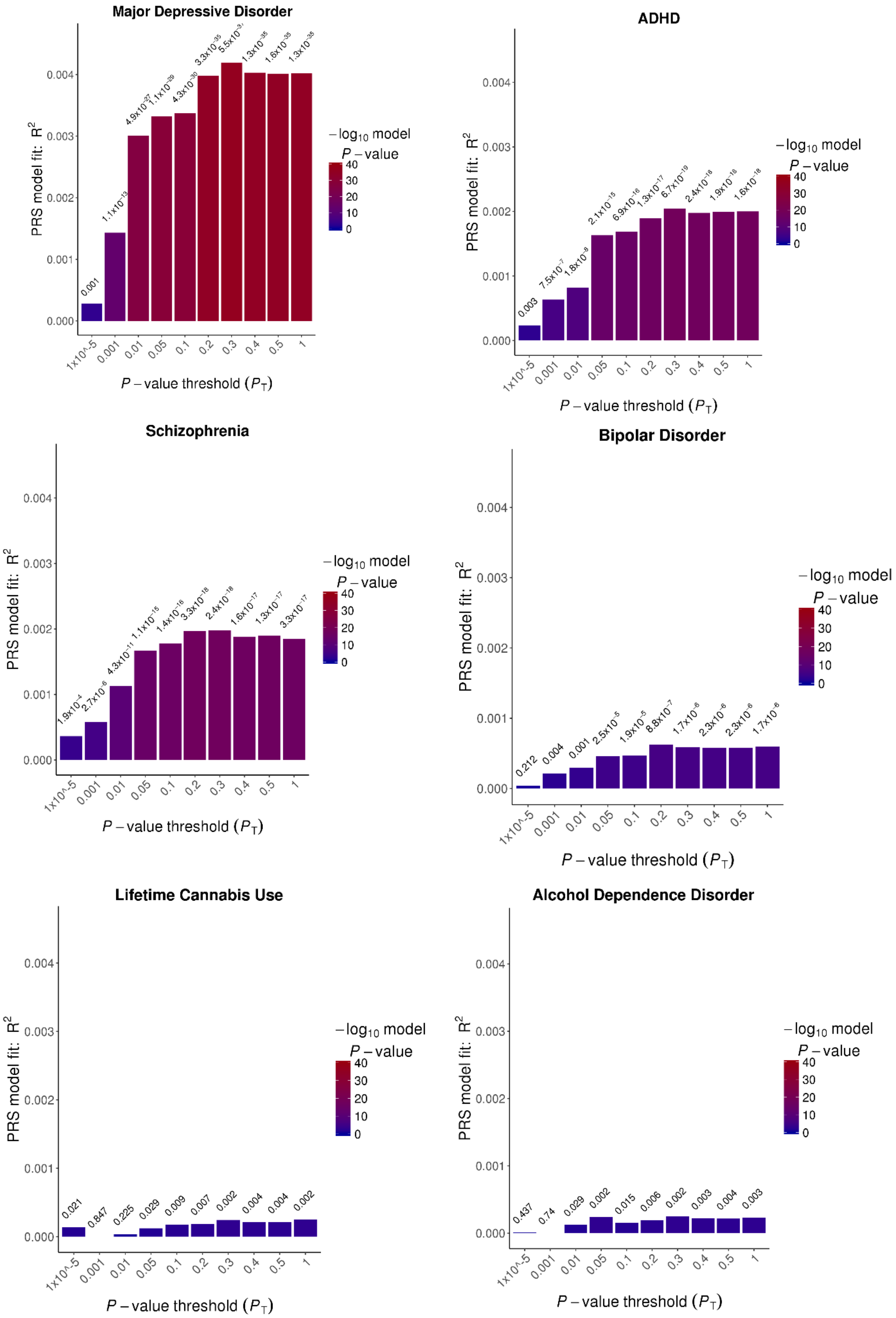

Figure S3. Pseudo $R^2$ plots of 6 PRS in predicting self-harm.
